# Novel visuomotor adaptation paradigm reveals a role of visual cortex in the plasticity of innate behaviors in mice

**DOI:** 10.64898/2026.04.17.719257

**Authors:** Ellery Jones, Massimo Scanziani

**Affiliations:** University of California, San Francisco; University of California San Francisco and Howard Hughes Medical Institute

## Abstract

A long-standing hypothesis in sensory neuroscience suggests that the evolutionary expansion of cortex in mammals may contribute to sensory-dependent adaptation by acting on subcortical pathways that drive innate behavior. However, direct experimental evidence is lacking. Taking the visual system as a model, it is known that there is significant interaction between the evolutionarily conserved Superior Colliculus (SC) and the comparatively modern Visual Cortex (VC), key structures in the mammalian visual system. In the SC, local alignment is established between a retinotopic map of the visual field and a map of orienting movement vectors during development and drives accurate visually guided orienting behavior throughout an organism’s lifespan. Interestingly mammals, like humans and non-human primates, readily adapt to altered visual experiences, while evolutionarily older vertebrates, like amphibians, lack this behavioral plasticity. To address this outstanding question, we have developed a novel behavioral paradigm for inducing visuomotor adaptation in freely moving mice that is analogous to paradigms utilized in primates. Our paradigm combines a visually guided orienting task and a novel mouse prism goggle system to shift the visual field. Using this paradigm, we demonstrate for the first time that mice gradually adapt to a chronic shift of their full visual field, suggesting this type of behavioral plasticity is conserved across mammalian species. Furthermore, we show that lesioning primary visual cortex (V1) prior to shifting the visual field disrupts normal visuomotor adaptation, suggesting that VC may play a generative role in the plasticity of fundamental visually guided behaviors. These findings lend support to the hypothesis that a particular evolutionary benefit of sensory cortex is the allowance for experience-dependent behavioral plasticity.

## Introduction

Sensorimotor transformations are the process by which sensory information is translated into an appropriate behavioral response. These transformations are the basis of fundamental behaviors like hunting for prey or fleeing towards shelter. Considering that sensory information (i.e. photons, soundwaves, vibrations etc.) and motor commands (neural signals for muscle contractions) occupy separate modalities the challenge of computing a motor command from sensory information is non-trivial, yet common to all animals. One way in which the brain solves this computation is through the alignment of anatomical maps of sensory and motor space. In fact, an evolutionarily ancient structure in the midbrain, the superior colliculus (SC) is a classic example. The SC is an evolutionarily conserved hub of visuomotor transformations ^1–4^ which drives visually guided orienting behaviors across vertebrates ^1,5–11^. The SC is anatomically organized such that the superficial layers contain a retinotopic map of visual space while the deep layers contain a map of movement vectors spatially aligned with the visual map ^12–14^. This local visuomotor alignment within SC underlies the accuracy of physical orienting movements towards a visual cue, (i.e. visually guided orienting behavior)^5,9,15,16^.

While sensorimotor alignment is defined early in development^17–19^, an organism’s sensory and motor experiences change throughout the course of its lifetime as it grows and matures. This creates a paradox in which hardwired sensorimotor alignment must also be plastic so that an organism can execute accurate sensory-guided behaviors throughout its lifespan. Neuroscientists have spent decades theorizing about how sensorimotor alignment at the neural circuit level adapts to maintain sensorimotor coordination at the behavioral level. In other words, how does the accuracy of goal-oriented movements recover when sensory information becomes unreliable? Taking the visual system as a model, it has been shown that humans and non-human primates readily adapt their visually guided behavior to a prism-induced visual field shift ^20–23^. In striking contrast, classic experiments by Roger Sperry demonstrate that amphibians permanently fail to orient towards their prey following an experimentally induced visual field shift^24^. Due to this difference in behavioral plasticity between vertebrate species, it has been suggested that the evolutionary expansion of the cortex in mammals underlies the ability to adapt to altered sensory experience. However, there remains limited direct evidence to support this decades old hypothesis.

The SC and the visual area of the cortex (VC) make up two distinct yet interconnected visual pathways. While these brain areas independently receive visual information from the retina, VC sends dense projections to SC^25,26^, creating an anatomical substrate by which VC may influence collicularly mediated behavior, such as accurately orienting towards a salient visual cue. <u>In fact, we hypothesize that VC contributes to the generation of behavioral plasticity via modulation of local visuomotor coordination in the SC.</u> To reveal the relative contributions of VC and SC to behavioral plasticity in mammals, we have developed a novel behavioral paradigm for inducing and quantifying visuomotor adaptation in freely moving mice that is analogous to paradigms utilized in primates. Our paradigm combines a visually guided orienting task and a novel mouse prism goggle system to shift the visual field. Using this paradigm, we demonstrate that adaptation to a chronic visual field shift is not specific to primates, rather is conserved across a wide subset of mammals. In addition, using circuit dissection techniques, we present data that support that, under normal visual conditions, the SC is sufficient for accurate visually guided behavior, but that the VC is necessary for the plasticity of these behaviors.

## Results

### A Novel Visually Guided Orienting Task (VGOT): expert mice learn to orient accurately to a visual cue

To characterize visuomotor adaptation in an ethological context, we designed a novel paradigm for inducing and quantifying visuomotor adaptation in freely moving mice (Fig. 1). We first designed a self-initiated <u>v</u>isually <u>g</u>uided <u>o</u>rienting task (VGOT) which allows mice to report the perceived location of a visual stimulus. The task is run under pitch dark conditions and mice are lightly food deprived prior to the start of behavioral training for the task. Mice trigger each trial by lining up at the “home port” facing the 0° position. Immediately after the trial initiates, there is a very brief (200ms) flash of light at the “target port” indicating reward availability at that location. Following trial initiation, mice approach the target location. If they arrive at the target within the 1 second response window, the target port dispenses a liquid reward. Once the mouse reaches the target, the trial ends and the mouse is able to return to the home port to initiate the next trial as the target port moves to a new random location within the arena. The physical target port moves between discreet locations across the azimuthal plane of the visual field (Fig. 1a) and the visual cue occupies 1.16° of visual space when the mouse is at the home port. Mice are considered to have reached expert criterion once they have met or exceeded experimenter defined thresholds of efficiency and accuracy for at least two consecutive sessions.

**Figure 1.**
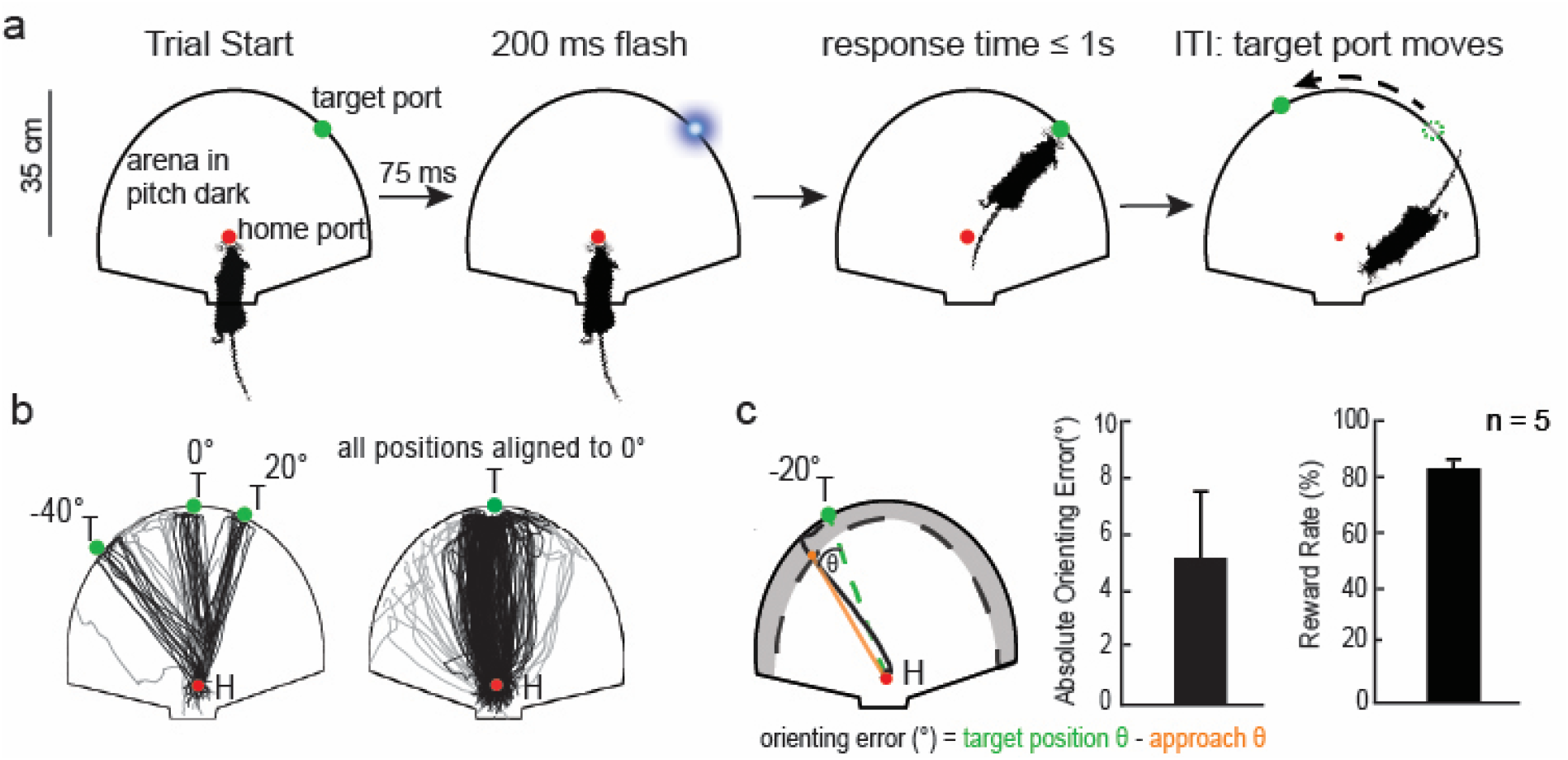
A novel paradigm quantifies visual spatial perception in freely moving mice. **A**. A novel visually guided orienting task (VGOT) allows mice to report their visual perception by orienting towards and approaching a brief flash of light in an otherwise pitch-dark arena. 0) The task 1) Mice initiate every trial by lining up at the home port facing the 0° position and a brief flash of light (cue duration = 200ms) is presented at the target position. 2) Mice have a 1000ms response window in which to approach the target. 3) Mice are rewarded for reaching the target location within the response window or a penalized with a 12s timeout if they fail to arrive at the target within the window. 4) In between trials the target randomly moves to a new location and the mouse returns to the home port to initiate the next trial. **B**. Raw approach traces of expert mice sorted by 3 example target positions and all trials from a session in which approach traces are rotated to align with an approach to the target at 0. **C**. Raw approach traces from the home port to the “edge zone” are used to define the approach orienting trajectory and compute the approach angle on a trial-by-trial basis. Approach accuracy is quantified by the orienting error (target position-approach). Expert mice complete the VGOT with high accuracy and efficiency (orienting error = 5.3° ± 2.5°, reward rate = 82.1% ± 5%). Absolute orienting error and reward rate plotted as mean ± sd.

Visualization of the mice’s raw approach paths to the target reveals that mice can reliably and accurately orient towards the target at various locations across the visual field (Fig. 1b, left). Aligning paths to all target locations to the 0° position, illustrates that mice directly approach the correct target location throughout a session (Fig. 1b, right). Approach accuracy and precision were quantified by computing the mice’s orienting error and reward rate for each behavioral session. The absolute, or directionless, orienting error is calculated for each trial by taking a linear projection of the approach path from the starting point (home port) to the point when the mouse first enters the “edge zone” (Fig. 1c, left). The projected approach trajectory is then used to compute the angle of the approach taken. The orienting error for that trial is the difference between the angle of the target position and the angle of approach. Once mice have reached “expert” criterion, they are able to complete the VGOT with an average orienting error of ~5° (5.3° ± 2.5°, Fig. 1c, center) with a reward rate around 80% (82.1% ± 5%, Fig. 1c, right). With this paradigm, we were able to quantify baseline mouse visual spatial perception while they freely performed an ethologically relevant task under normal visual conditions.

### Novel Mouse Goggle Systems induces full visual field shift

With baseline orienting behavior established on the VGOT, we created a novel mouse goggle system to induce visuomotor adaptation in freely moving mice (Fig 2a,b). The mouse goggle system consists of two components: a surgically implanted goggle base and clip-on lens frames (Fig. 2a). Both the goggle base and lenses are lightweight (base = 1.2g, lens frames = 0.8g) to minimize disruption in the mice’s ability to move freely. The goggle bases are implanted prior to the start of experiments, and, after recovering from surgery, mice live and behave normally with the base. The lens frames clip-on to the goggle bases using tiny magnets (Fig. 2a, right panel). Control lenses are constructed by covering lens frames with clear plastic to create a normal visual experience and habituate mice to wearing goggle lenses. Prism lenses are constructed by attaching Fresnel prisms to the lens frames (Fig. 2b, left panel). Prism lenses shift the visual field along the azimuthal plane such that the perceived location of an object directly in front of the viewer is offset by 40° (38 ± 1.8°, Fig. 2b, center and right panels). The clip-on lens system allows the experimenter to easily interchange between maintaining normal visual experience with the control lenses or to chronically, yet reversibly, shift the mouse’s full visual field using the prism lenses.

**Figure 2.**
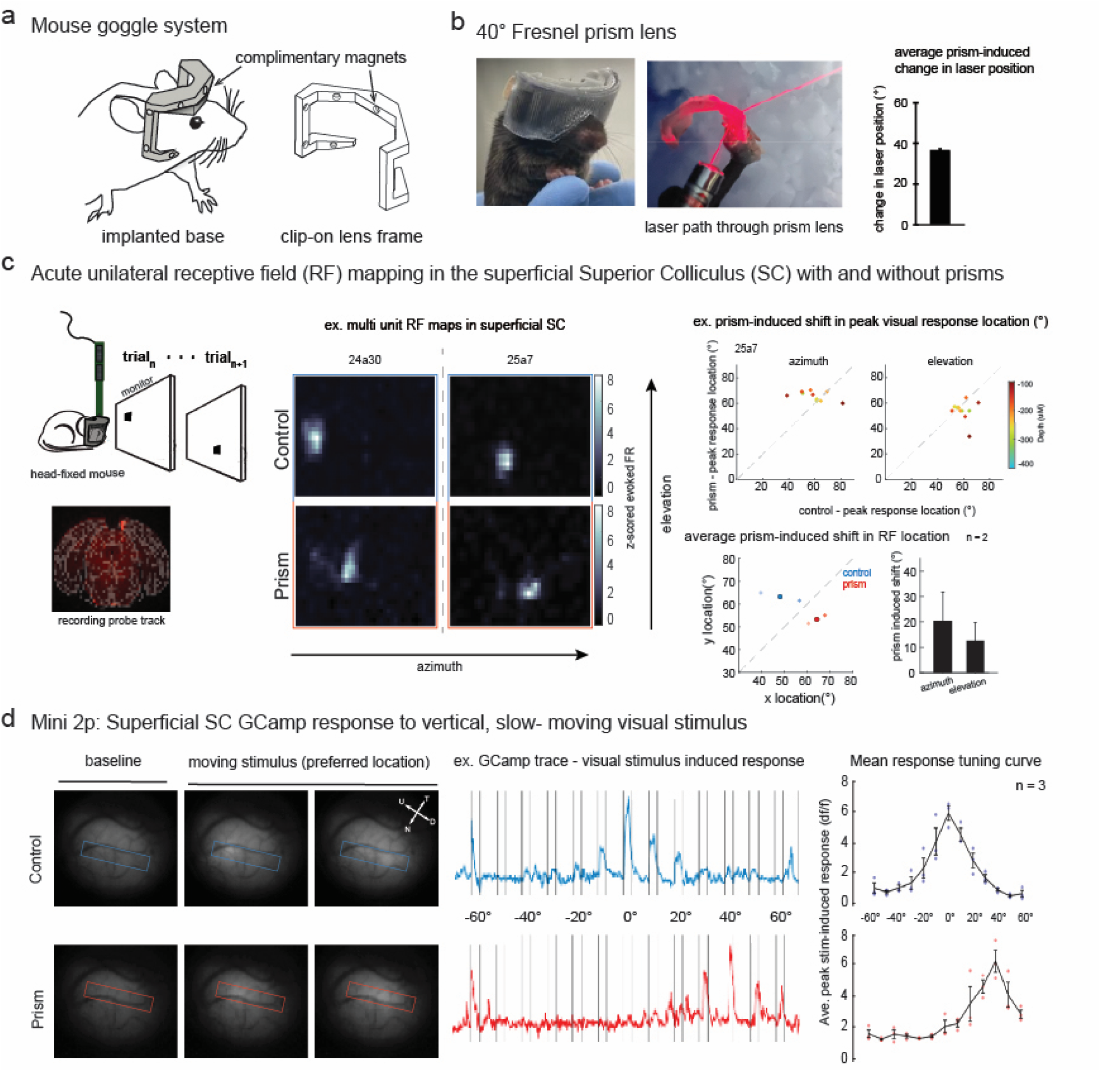
Novel mouse goggle system induces full visual field shift. **A**. A novel two-part mouse goggle system. Goggle bases are surgically fixed to the mice’s skulls and lens frames clip-on to the bases with small magnets. **B**. Control lenses provide normal visual experience while prism lenses use Fresnel prisms to create a 40° horizontal shift in the visual field. Prism lenses generate a near 40° (38 ± 1.8°) deflection in laser light paths **C**. Mouse prism goggles induce a visual field shift at the level of the visually responsive cells in the SC. Acute, head-fixed, unilateral electrophysiological recordings were performed in the right hemisphere of SC while mapping the location of the maximal stimulus evoked response (RFs) under normal visual experience and with prism lenses. RF location shifts significantly along the azimuth when mice are wearing prism lenses but is relatively stable across elevation within and across animals (*Δ* peak location _control-prism, azimuth_ = 19.26 ± 12.93°, p _azimuth_ = 0.002, *Δ* peak location _control-prism, elevation_ = 11.15 ± 9.85°, p _elevation_ = 0.068) **D**. Using UCLA miniscope calcium imaging (GCaMP8m), calcium responses of cells in superficial SC were recorded as a black square moved vertically down the screen at various locations across the azimuthal plane. Typical visual responses were recorded during normal visual experience and with prism lenses. The location within the superficial SC of the region of interest with the highest intensity response (ROI_peak_) shifted anteriorly and laterally with the prism lenses relative to when visual experience was normal. Plotting the calcium traces in the ROI_peak_ of individual mice reveals a peak visual response when the stimulus is at 0° during normal visual experience while the ROI_peak_ with prism lenses showed a maximal response when the stimulus is at 40°. This prism-induced shift in preferred stimulus location for the ROI_peak_ was consistent across animals.

To validate that the mouse prism lenses shift the visual field within the mouse visual system, we mapped the locations of the visual receptive fields (RFs) of visually responsive cells in the superficial layers of the SC (Fig. 2c). A visual RF defines the area in visual space in which a stimulus evokes the maximal electrophysiological response from a cell, i.e. the cells preferred stimulus location. To map visual RFs in SC, we presented head-fixed mice with a high contrast sparse noise stimulus while unilaterally recording across the superficial layers of SC with a linear probe (Fig. 2c, left panel). For each trial, a black rectangle that occupies 10° of visual space was briefly presented (stimulus duration = 100ms) at random locations across the monitor. Over the course of mapping, each stimulus was presented 30 times at each location, and the RF is visualized by plotting a heat map of the z-scored, stimulus-evoked firing rate at each stimulus location across all trials. To visualize and quantify the prism goggle induced a 40° visual field shift, visual RFs were mapped in two consecutive conditions. First, visually evoked responses in SC were recorded under normal visual experience to locate the innate RF location of the cells being recorded (Fig. 2c, center panel, top row). Immediately following the control recording, and without moving the probe, mice are fit with prism lenses and RF mapping is repeated, revealing a shift in RF location primarily along the azimuth (Fig. 2c, center panel, bottom row). Within animals, across the superficial layers of SC, the location of peak visual responses with prism lenses shifted relative to the peak location during normal visual experience primarily along the azimuth with relatively stable locations across elevation (Fig. 2c, center panel, bottom row). On average, across animals, the RF location shifted significantly only along the azimuth(*Δ* peak location_control-prism, azimuth_ = 19.26 ± 12.93°, p _azimuth_ = 0.002, *Δ* peak location _control-prism, elevation_ = 11.15 ± 9.85°, p _elevation_ = 0.068).

Visually responsive cells in SC are known to respond strongly to small, moving stimuli ^27^. Using the <u>UCLA</u>Miniscope and a calcium indicator (GCaMP8m), we recorded calcium responses of cells in superficial SC to a black square moving vertically down the screen at various locations across the azimuth. Similarly to the acute physiological recordings, typical visual responses were recorded during normal visual experience (Fig. 2d, left panel, top row), and responses with prism lenses were recorded immediately following (Fig. 2d, left panel, bottom row). The location within the superficial SC of the region of interest with the highest intensity response (ROI_peak_) shifted anteriorly and laterally with the prism lenses relative to when visual experience was normal (Fig. 2d, left panel). The direction of this shift is consistent with the known retinotopy of the mouse SC, in which anterolateral cells respond to temporal areas of the visual field ^28^.

Plotting the calcium traces in the ROI_peak_ of individual mice reveals a peak visual response when the stimulus is at 0° during normal visual experience (Fig. 2d, center panel, top row) while the ROI_peak_ with prism lenses showed a maximal response when the stimulus is at 40° (Fig. 2d, center panel, bottom row). This prism-induced shift in preferred stimulus location for the ROI_peak_ was consistent across animals (Fig. 2d, right panel). These data show that, at the sensory level, the prism goggles induce an instantaneous shift in the full visual field of a mouse.

### Mice gradually adapt to a chronic visual field shift

Combining the VGOT and mouse goggle system, we have been able to longitudinally measure visuomotor adaptation in freely behaving mice (Fig. 3a). Once baseline orienting behavior is established in the absence of any lenses, control lenses are placed to generate a baseline which accounts for any effect of wearing the lenses themselves. Even with control lenses, mice maintain their baseline orienting accuracy (Fig. 3b). The clip-on design of the goggle lenses allowed us to start the mice in a session with control lenses for 75-100 trials and then remove the mouse from the task and swap control lenses for prism lenses under complete darkness.

**Figure 3.**
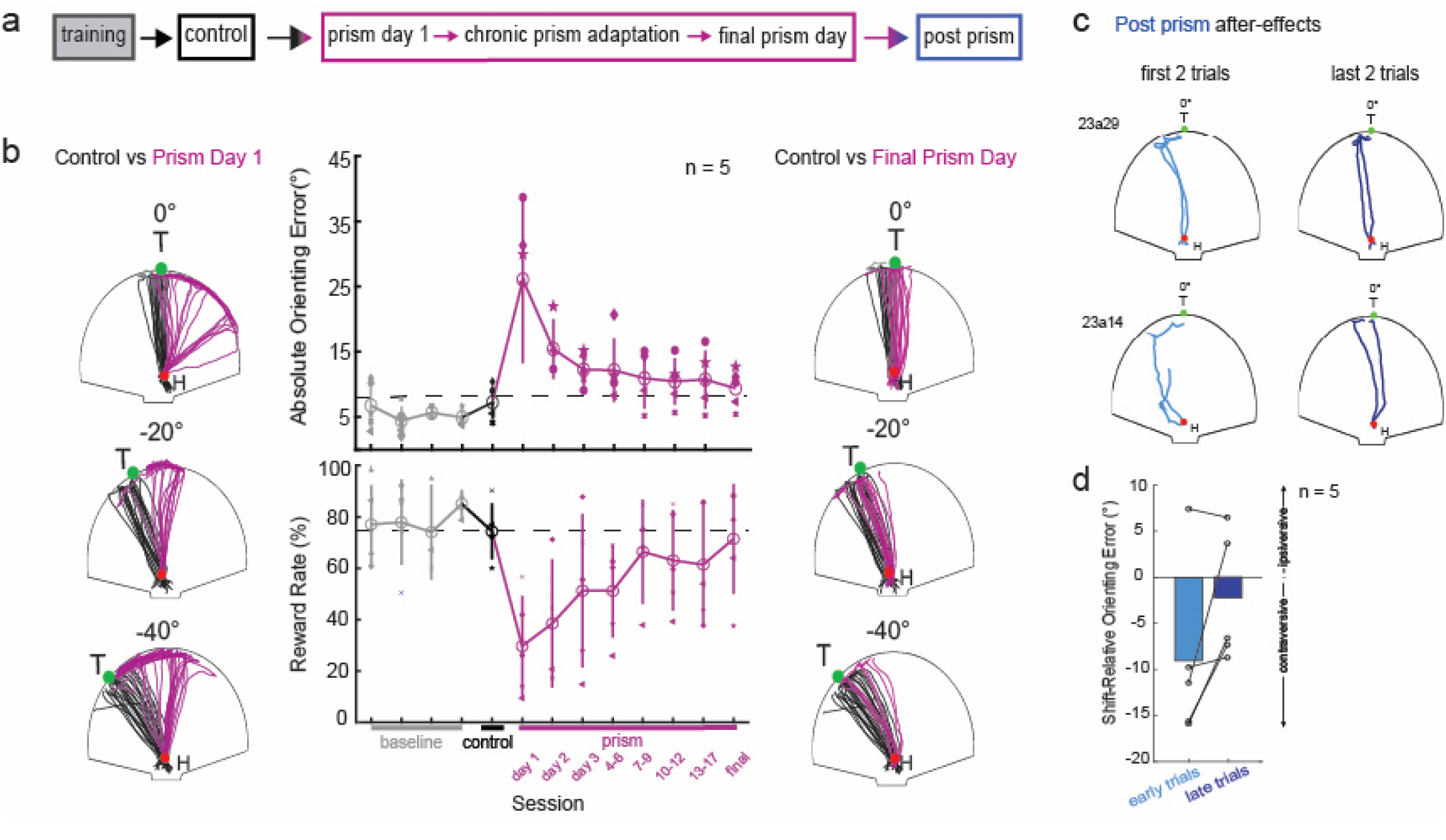
Mice gradually adapt to a prism-induced full visual field shift. **A**. Protocol for measuring behavioral adaptation to a chronic, yet reversible prism induced visual field shift in mice. Note that the sessions transitioning from one condition to another occur consecutively on the same day (i.e. the final control session to first prism session, and final prism session to first post-prism session). **B**. Comparison between raw approach traces in control conditions and immediately after prism-induced visual field shift exemplifies instantaneous shift in approach behavior. Baseline orienting error refers to orienting accuracy specifically during the final training sessions, without any lenses. Control orienting error reflects the orienting accuracy during the final sessions with control lenses. Demonstrates the lack of disruption caused specifically by the addition of lenses. The prism induced shift in approach behavior is reflected by large increase in orienting error during the first behavior session with prism lenses. Orienting error gradually improves over the course of chronic visual field shift until returning to baseline orienting accuracy after 18 ± 3 sessions (orienting error _prism day 1_ = 26 ± 15°, orienting error _final prism day_ = 9.5 ± 4°). The majority of adaptation occurs within the first 4-6 sessions (orienting error _first half_ = 25 ± 12.5°, orienting error _second half_ = 2.5 ± 5°)). Comprehensive adaptation is exemplified by comparing raw approach traces under control conditions to raw approach traces on the final day of prism wear. **C**. Approach paths for the first and last two trials during the post-prism behavior session demonstrate the after-effects which appear transiently following restoration of normal visual experience. **D**. On average, mice make larger, contraversive orienting errors during early trials following prism removal but have regained baseline orienting accuracy by the end of the behavioral session.

The mouse is then immediately returned to the VGOT arena to begin the first prism session (Fig. 3a). Placing the prism goggles causes an immediate shift in orienting behavior which is consistent with the direction and magnitude of the visual field shift (Fig. 3b, left panel approach traces) and reflected in the significant increase in orienting error during the first session with prism goggles (prism day 1, orienting error = 26 ± 15°, Fig. 3b, center panel, top).

Over the course of ~2.5 weeks (18 ± 3 days) of chronic wear of the prism lenses, the average orienting error across VGOT sessions gradually returns to baseline level orienting errors (final prism day, orienting error = 9.5 ± 4°, Fig. 3b, center panel, top). Across animals, the majority of adaptation occurred within the first six days of the chronic visual field shift (*Δ* orienting error _first half_ = 25 ± 12.5°, *Δ* orienting error _second half_ = 2.5 ± 5°). Visual comparison of raw approach paths to the target under control conditions and on the final day of prism-wear illustrate the qualitative similarity of the approach trajectories before and at the conclusion of the adaptation period (Fig. 3b, right panel). Consistent with the changes in orienting accuracy over the course of adaptation, the average reward rate per session falls to chance levels upon placement of the prism goggles and recovers to baseline rates by the end of the adaptation period (Fig. 3b, center panel, bottom).

Within the field of primate prism adaptation, true visuomotor adaptation is thought to be marked by motor errors in the opposite direction of the original visual field shift once prisms are removed (i.e. after-effects) ^29^. The presence of after-effects is immediate but dissipate rapidly. Immediately following the final VGOT session with prism lenses, prisms were replaced with control lenses and the mice were returned to the VGOT. Early in the post-prism session, approach paths reveal orienting errors in the opposite direction of the prism induced visual field shift (Fig. 3c, left). However, by the end of the post prism behavior session, mice take direct approaches to the target with near baseline accuracy (Fig. 3c, right). On average mice’s orienting error exhibit after-effects which are gone by the end of the behavior session (orienting error_early trials_ = 9.7 ± 8.2°, orienting error _late trials_ = 2.8 ± 7.1°, Fig. 3d). Taken together, these data demonstrate, for the first time, that mice are capable of adapting to a chronic disruption in visual experience.

### Primary Visual Cortex (V1) is not necessary for the execution of learned visually guided behavior

Having established what visuomotor adaptation looks like in freely moving mice under normal circumstances, we then endeavored to uncover the necessity of primary visual cortex (V1) for the plasticity of visually guided orienting behavior. To do so, we first needed to address the assumption that mice’s performance on the VGOT is driven by the visuomotor coordination of the superior colliculus (SC). If this is the case, then V1 would not be necessary for execution of the task once it has been learned. To test this hypothesis, we trained a cohort of mice on the VGOT task until they reached expert criterion and quantified their baseline performance with control lenses (Fig. 4a). We then bilaterally removed V1 and allowed mice to recover from surgery before retesting their orienting accuracy in the absence of V1, but with normal visual experience, prior to starting prism adaptation (Fig. 4b). Comparing coronal sections of the brains of mice with V1 lesions with equivalent sections of the Allen Mouse Brain Atlas^30^ demonstrate that the lesions are largely restricted to V1 and the higher visual areas (HVAs) are spared (Fig. 4b). This allowed us to evaluate the necessity of V1 for the execution of visually guided orienting behavior in expert mice. Prior to V1 being lesioned, mice performed at expert criterion under control conditions (orienting error = 6.64 ± 2.11°, reward rate = 77.4 ± 13.8%, n = 5, Fig. 4c). In the immediate sessions following bilateral removal of V1 and recovery from surgery, accuracy and efficiency on the VGOT decreased slightly (orienting error = 15.24 ± 5.31°, reward rate = 56.2 ± 22.2%), but returned to baseline behaviors after a few control behavior sessions with normal visual experience (orienting error = 4.88 ± 0.91°, reward rate = 92.1 ± 1.7%, Fig. 4c). These data demonstrate that, in mice, V1 is not necessary for the execution of visually guided orienting behavior under normal conditions and are consistent with differential roles in visual processing typically ascribed to SC and VC (i.e. visual detection and orienting behavior, or visual discrimination respectively). Given these data, we proceeded to evaluate if V1 may be specifically involved in the plasticity of visually guided behavior.

**Figure 4.**
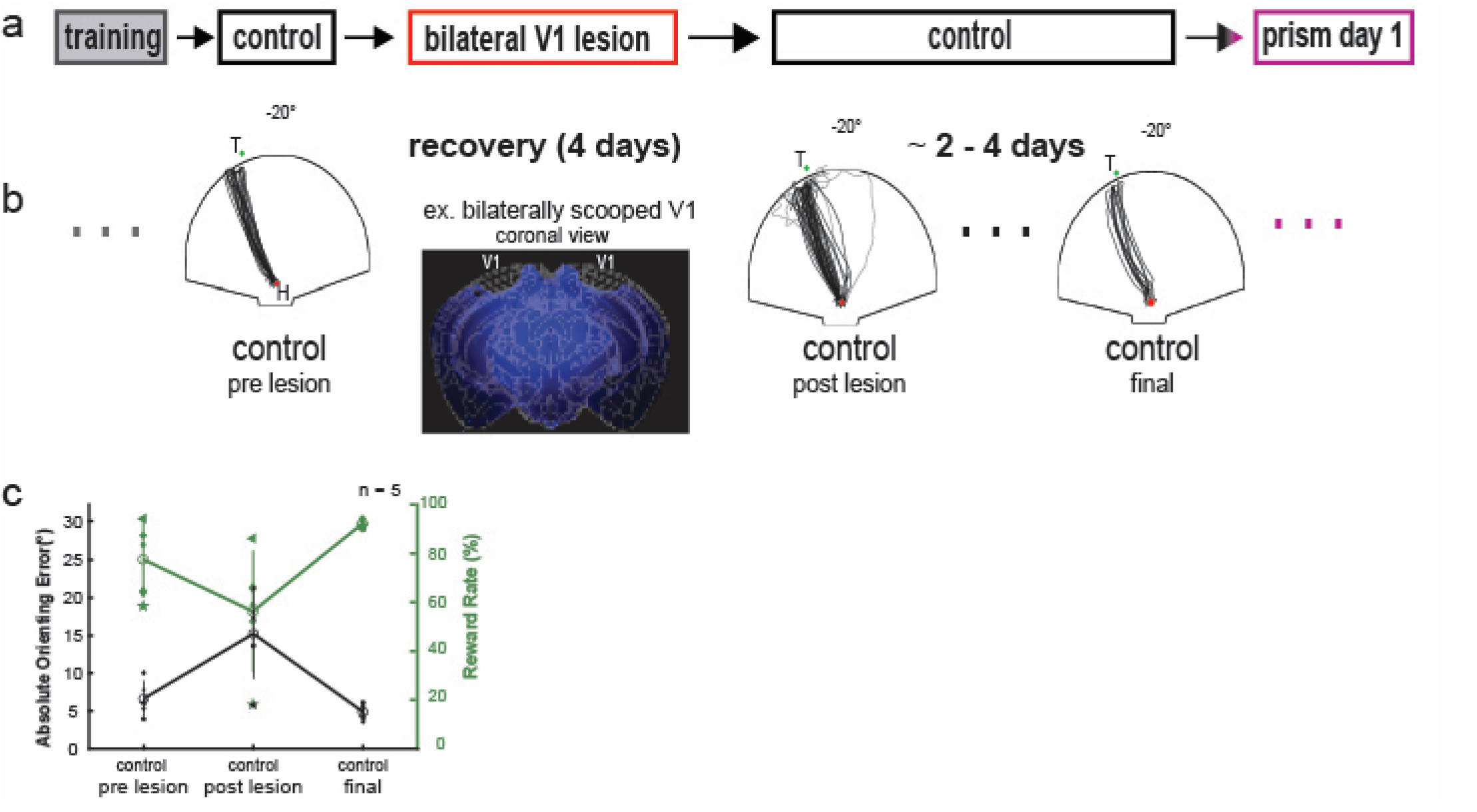
Primary visual cortex is not necessary for the execution of established visually guided orienting behavior. **A**. Protocol for testing visuomotor adaptation in mice with bilateral V1 lesions. Mice are trained to criterion before quantifying baseline orienting accuracy during normal visual experience. V1 is removed bilaterally, and mice return to the VGOT under control conditions after recovering from surgery. Once baseline orienting accuracy is re-established post lesions, mice are fit with prism goggles and adaptation is measured. **B**. Visualization of example lesions via coronal sectioning with DAPI staining and optical laser surface imaging. Exemplar raw approach paths before and after lesioning V1. The middle panel is immediately after the surgical recovery period, while the last panel is from the final session before prism adaptation begins (typically 2-3 behavior sessions post lesion recovery). **C**. Lesioning V1 does not impact the execution of visually guided orienting or performance of the VGOT task once it is already learned. Orienting accuracy and reward rate worsen slightly in the VGOT sessions immediately following lesions but quickly return to baseline (orienting error _pre lesion_ = 6.64 ± 2.11°, reward rate _pre lesion_ = 77.4 ± 13.8%, orienting error _post lesion_ = 15.24 ± 5.31°, reward rate _post lesion_ = 56.2 ± 22.2%, orienting error _final control_ = 4.88 ± 0.91°, reward rate _final control_ = 92.1 ± 1.7%).

### Loss of V1 disrupts normal visuomotor adaptation in adult mice

Immediately following the final VGOT session with control lenses, mice lacking V1 were fit with prism lenses that induced a 40° horizontal shift in the visual field. Similarly to what was observed in intact animals, prism lenses caused an immediate shift in orienting accuracy, reflected by a nearly 6-fold increase in orienting error and a drop to chance levels in reward rate on prism day 1 (Fig. 5a, left panel, *mean orienting error = 30°, mean reward rate = 50%, n = 5*). However, without V1, mice have a much slower adaptation rate to a chronic, prism-induced visual field shift compared to intact animals (Fig. 5a, center panel, top) and fail to recover baseline orienting accuracy in the same number of sessions (Fig. 5a, left panel,Δ *mean orienting error final prism day – baseline: lesioned mice = 10°, intact mice = 2*.*5°*). Interestingly, the effect of lesioning V1 seems to impact the adaptation rate most severely within the first four to six days of chronic prism-wear (i.e. the initial adaptation period) (Fig. 5a, center panel, prism day 1 to prism day 6 *slope: lesioned mice = -1*.*7, intact mice = -4*.*3*). Optical laser imaging (Fig. 5b) was used to quantify the extent of each lesion. Total cortical tissue removed was relatively consistent across animals, with the spread of the lesions being most consistent (lesion perimeter: 7.41 ± 1.08 mm). We found a strong correlation between adaptation rate and the spread of the lesion during the initial adaptation period (R _perimeter_ = 0.965, p _perimeter_ = 0.008, Fig. 5c, top). These data suggest that loss of V1 is most disruptive to visuomotor adaptation immediately following the prism-induced visual field shift. The lateralization of visual processing streams makes it possible that the impact of V1 lesions may differ based on whether it is ipsilateral or contralateral to the direction of the prism induced visual field shift. The significance of the strong correlation between the initial adaptation rate and spread of the lesion seems to be driven by the spread of the lesion contralateral to the direction of the visual field shift (R _perimeter, contra_ = 0.924, p _perimeter, contra_ = 0.025, R _perimeter, ipsi_ = 0.876, p _perimeter, ipsi_ = 0.051). These data suggest that V1, and especially the hemisphere contralateral to the visual field shift, is involved, particularly in the early phases of behavioral adaptation to a chronic visual field shift.

**Figure 5.**
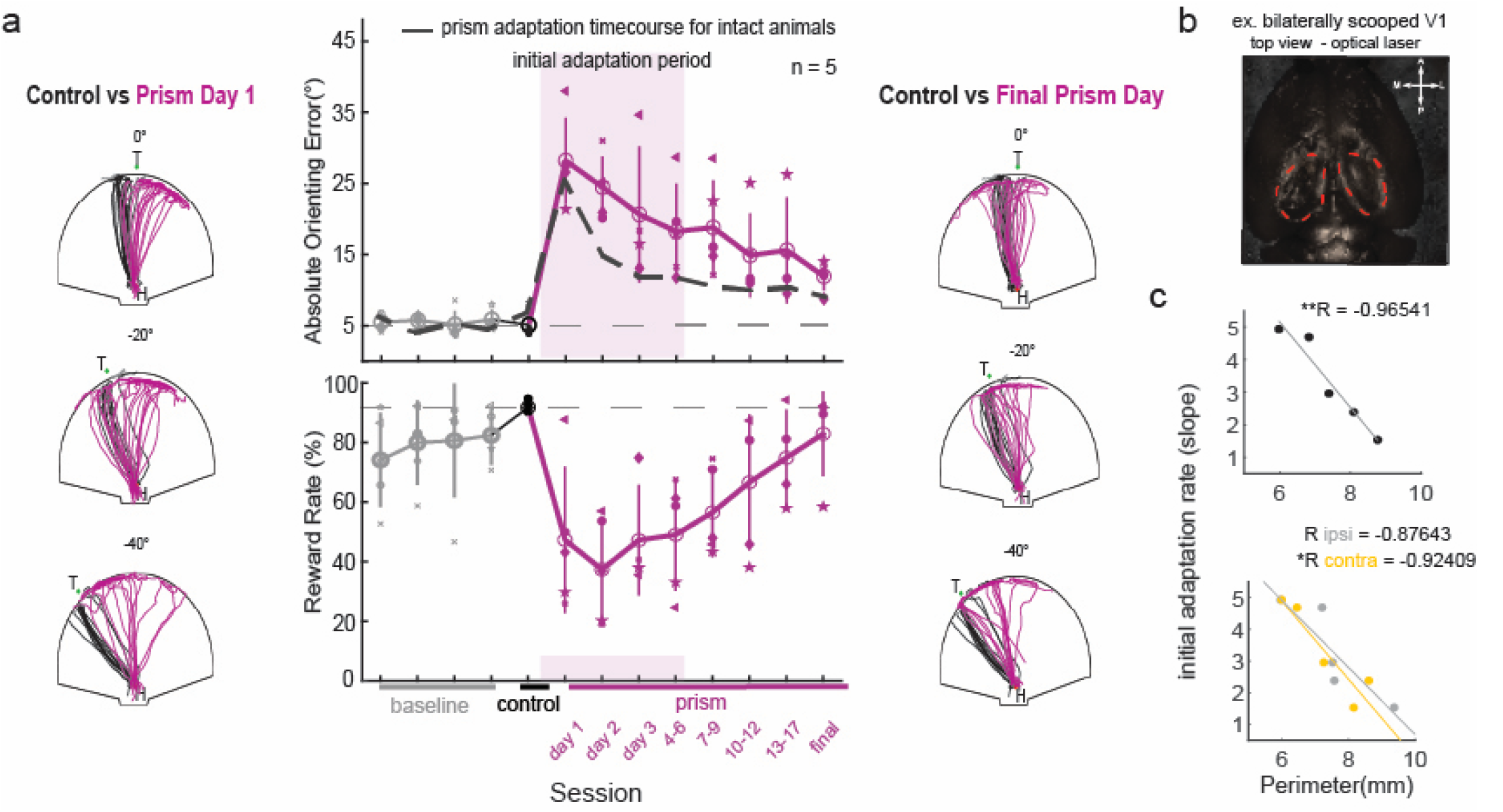
Primary visual cortex plays a potentially generative role in the plasticity of visually guided behavior. **A**. Lesioning V1 disrupts the rate of visuomotor adaptation compared to intact animals. Comparison of raw approach traces during the final control session and the first session with prism lenses in animals with bilateral V1 lesions illustrates expected prism-induced shift in orienting behavior. In contrast to intact animals (dashed-line), lesioned mice display delayed adaptation in both rate and magnitude, particularly during the initial adaptation period (Δ mean orienting error final prism day – baseline: lesioned mice = 10°, intact mice = 2.5*°*, initial adaptation rate: lesioned mice = -1.7, intact mice = -4.3). Raw approach traces on the final day of prism wear are notably different from approach traces under control conditions. **B**. Optical laser imaging was used to quantify the extent of each lesion. **C**. Between group differences in initial adaptation rate (deg/session) is driven by the spread of V1 lesions in the hemisphere contralateral to the direction of visual field shift. Dimensions related to the spread of lesions were very highly correlated with the adaptation rate during the initial adaptation period (R _perimeter_ = 0.965, p _perimeter_ = 0.008). The correlation between lesion spread and the initial adaptation rate is driven by the spread of lesions in the hemisphere contralateral to the direction of the visual field shift (R _perimeter, contra_ = 0.924, p _perimeter, contra_ = 0.025, R _perimeter, ipsi_ = 0.876, p _perimeter, ipsi_ = 0.051).

## Discussion

Quantification of visuomotor adaptation in a variety of vertebrate species has highlighted the difference in behavioral plasticity in species with a cortex (i.e. mammals) and without (i.e. reptiles, birds), leading to a decades-long hypothesis that an evolutionary advantage of cortex is experience-dependent optimization of behavior. It has specifically been shown that animals which evolved a more expansive/complex cerebral cortex (i.e. primates, cats) readily adapt their visually guided behavior to a chronic, prism-induced visual field shift. The present study uses mice, a readily available animal model, to investigate this long-standing hypothesis at the behavioral and neural circuit level. To do so, we first established a novel paradigm that induces visuomotor adaptation in a manner similar to prism adaptation paradigms classically used for humans and non-human primates. We demonstrate that freely moving mice can be trained to perform a new ethologically relevant visually guided orienting task that simulates a prey-capture like behavior. Once reaching expert levels, mice use vision to perform this task with high accuracy, despite concerns regarding lower visual acuity of mice. We introduce a novel mouse goggle system which allows for chronically yet reversible manipulation of mouse visual experience without disrupting normal visually guided behavior. With this system we show that putting on prism lenses causes an instantaneous shift in the visual field, at both the level of visually evoked neural activity and visually guided orienting behavior. The direction and magnitude of the shift correspond with the direction of the prism-induced shift. Our results demonstrate, for the first time, that mice are capable of adapting to a significant alteration of their visual experience, a phenomenon that has previously only been reported in human and non-human primates ^20-23,31^.

This novel paradigm additionally provides a platform for investigating neuroplasticity at the behavioral and circuit level across a wide range of contexts (e.g. development, aging, health and disease). Combining an ethologically relevant, freely moving behavior with clear goals and tight experimental control of sensory experience, opens the door for resolving lingering questions about the mechanisms of experience dependent adaptation, a process fundamental for all organisms to mature and survive in a dynamic environment.

Having described what visuomotor adaptation typically looks like, we systematically tested the contribution of V1 to the execution and plasticity of established visually guided orienting behavior. We demonstrate that V1 is not actively necessary for the execution of visually guided behaviors established prior to the cortical lesion. This result is consistent with our assumption that the visually guided orienting task is mediated by visuomotor coordination within the SC. Importantly, we show for the first time that without V1, mice failed to adapt normally to the chronic visual field shift. All together the data presented in this study provide direct evidence supporting the hypothesis that the metabolically costly evolutionary expansion of the cerebral cortex was offset by the addition of increased behavioral plasticity to cope with a dynamic environment through the modification of subcortical circuits.

In addition to superficial SC, V1 projects to numerous targets and so it is worth specifically ablating the collicular projecting neurons in V1 to confirm that this is the direct pathway by which V1 might be influencing visuomotor alignment in the SC. Additionally, lesioning V1 did not completely abolish visuomotor adaptation, rather it significantly slowed and prolonged adaptation. This sparing of behavioral plasticity could be related to the fact that V1 relays visual information to higher visual areas (HVAs). HVAs also send anatomically distinct projections into the intermediate layers of SC, and as such could be the visual cortical regions driving visuomotor adaptation. Alternatively, it is not out of the question that SC in mammals has evolved independent mechanisms of adaptation, which is a line of inquiry that would need to be pursued further. However, our results are consistent with the findings that in ferrets, adaptation to monocular occlusion, as measured by performance on a sound localization task, is dependent on corticocollicular projections from primary auditory cortex to inferior colliculus ^32-33^. The similarity between these findings and our data in the visual corticocollicular system is consistent with the modular hypothesis of the organization of cortical circuits across sensory modalities. The results of this study provide a tractable yet ethologically relevant paradigm for studying learning and sensorimotor alignment across states of development, health and disease. We demonstrate here how this paradigm can be used to systematically probe outstanding questions of neural circuit computations. Our findings suggest that having a cortex may allow organisms the ability to update their actions in an experience dependent way in order to execute increasingly complex behaviors.

## Materials and Methods

### Visually Guided Orienting Task

#### Construction

*Hardware* - Behavior enclosures were built with the dimensions of 70cm long x 52.5cm wide x 90cm high using 25mm construction rails (ThorLabs) and 61cm × 61cm black hardboard (ThorLabs). The arena floor was custom cut from white waterproof high-density polyethylene (King Starboard^®^ Marine HDPE, TAP plastics) into a semi-circular shape (1.27cm thick x 76.2cm diameter) and mounted onto two rectangular panels (76.2cm long x 6.35cm wide x 6.35 cm high) arranged in an L shape to create a “false floor”. One end of a custom milled 38cm long x 0.5cm aluminum arm was attached to a step motor (model CTP20NLF17, Kollmorgen) and the other end was attached to a custom designed, 3D printed target port enclosure (Fusion 360, Autodesk, Formlabs). The step motor with the aluminum arm and target port was secured underneath the center of the arena floor, such that the target port sat 3.5cm above the arena floor. The top of target port enclosure housed a 5mm infrared (IR) LED facing the ceiling for target tracking and the front housed a 5mm standard LED (Mouser Electronics) and spout (16-gauge blunt tip syringe needle, Shintop) for visual stimulus and reward delivery respectively. A ~2mm hole was drilled in the arena floor, 35cm from the floor edge, to hold the spout for the home port. The arena walls were constructed from 1.5mm thick black plastic (Styrene High Impact Plastic, TAP plastics). The gap between the arena walls and floor edge, was filled with complementary strips of brush weather stripping (Fowong) to allow for the target to move smoothly along the circumference of the arena floor. To track animal behavior, a camera was mounted at the top of the enclosure (acA1300-200um, Basler) and the arena was illuminated by two custom built IR LED (930uM) arrays. A small speaker (FE83NV 3” Full Range Speaker, Fostex) was mounted to the top of the arena to mask any spatially relevant noise from the motor. Rewards were delivered using miniature solenoid valves (LHDA0533115H, Lee company). Microcontrollers (Arduino UNO and MEGA Rev3) communicated between the computer and the physical components within the arena, allowing for closed loop control of the task. *Software –* Visually reactive programming was used to achieve closed-loop control of the task. Custom code utilized live pose estimation (Bonsai, DeepLabCut) to track the animal and trigger the various parameters of the task structure through communication with task hardware described above. User input determines the range of positions that the target will visit. The task code was written to allow for live acquisition of data about metrics of the animals’ performance.

#### Training

Mice were trained on the VGOT in two stages. 1) During the “Hallway” training stage, transparent dividers (TAP plastic) are used to limit access to the center of the arena and a range of +-20° along the azimuth. At this stage, mice are rewarded at both the home port, following trial initiation, and the target port and begin with an initial response window of 7 seconds. As they successfully complete trials, the reward at the home port is titrated out, while the target reward volume increases to its final volume. Concurrently, the response window progressively decreases to 1 second as the mice learn the foundational parameters of the task. Once mice are able to complete this phase of training with at least 75% reward rate, they graduate to the full phase of training. 2) The “Full” training phase is functionally equivalent to the full task. The transition to the “Full” phase is the first time mice are exposed to the entire arena and the full range of possible target locations. Mice stay in the “Full” training stage until they reach expert mice criterion of at least 75% reward rate and less than 10° orienting error.

#### Testing

After the completion of the final training session, mice are fit with control lenses to allow them to acclimate to wearing lenses overnight. Task structure is unchanged from the end of training. Mice repeat control sessions until they have a reward rate of at least 75% for two consecutive sessions. A subset of mice underwent control sessions with blank trials, during which the target cue light would not turn on after trial initiation during random trials. On the day that mice first switch to prism lenses, all animals start with their final control session. Mice are placed into the VGOT and allowed to complete at least 75 trials with a minimum 75% reward rate. Once the control session is completed, all light sources are turned off prior to removing the mouse from the arena. The control lenses are then switched for prism lenses and mice are immediately returned to the VGOT task. Mice wear prism lenses chronically. They are tested in the VGOT consecutively for the first three days of prism wear and then every other day thereafter until animals regain baseline orienting behavior. Once mice have adapted, they complete the final prism session, and then prism lenses are switched back to control lenses in a completely dark room. Mice are then returned with control lenses to the VGOT enclosure for a “post prism” session.

#### Read out

DeepLabCut based live pose estimation is used via Bonsai to track animals’ approach behavior throughout the entire prism session. Timestamps are taken for each major occurrence in the task structure (e.g. trial start, target approach times, rewarded approaches, etc.). Pose estimation and timestamp data are parsed post hoc to isolate approach behavior on a trial-by-trial basis. Raw approach paths are transformed into orienting trajectories that are used to calculate the orienting angle for each trial.

#### Mouse prism goggle system

##### Goggle construction

Both components of the goggle system (i.e. goggle base and lens frames) were 3D printed in house (Formlabs) from custom made design files (Fusion 360, Autodesk). Miniature magnets (D032-063, AmazingMagnets) were glued (Krazy glue, super glue) into 9 slots along the outward surface of the goggle bases and inward surface of the lens frames. Either custom cut clear acetate (TAP plastic) or 40 diopter Fresnel prisms (3M) were adhered to the surface of the lens frames to create control or prism lens respectively. *<u>Goggle base implant</u>*: Mice were anesthetized with 1.5-2% isoflurane and placed in a stereotactic set up (Kopf). Body temperature was monitored using a rectal probe and maintained at 37°C using a heating pad (FHC; DC Temperature Controller). Throughout the procedure, the animals’ eyes were protected by a thin layer of eye ointment (Bausch & Lomb). Animals fur on top of the head was shaved and skin was sterilized with Betadine before being removed to expose the skull.

Lidocaine (7 mg/kg) was injected locally at the site of incision. Two 3/32” bone screws were inserted into burr holes equidistant from bregma (-100uM anterior-posterior, +/-450uM medio-lateral) to serve as anchors. Goggle bases are then cemented to the skull using self-adhesive resin cement (RelyX Unicem2 Automix, 3M) and the entire surface was sealed with dental cement (Ortho-Jet acrylic resin, Lang Dental). At the end of the procedure, animals were administered a subcutaneous injection of 0.1mg/kg Buprenorphine as a postoperative analgesic.

#### Receptive field mapping in the superior colliculus (SC)

##### Goggle base implant and craniotomy

Prior to electrophysiological recordings, goggle bases were implanted as described above. While still under anesthesia, a craniotomy was made on the right hemisphere over SC (diameter ~400uM, +400uM anterior-posterior, +775uM medio-lateral) and protected by a local application of Kwik-Cast (WPI) until the day of recording. A custom designed head bar for head fixation was attached to the center of the base using resin cement (RelyX Unicem2 Automix, 3M).

##### Receptive field mapping

Visual stimuli were generated in Matlab with Psychtoolbox and custom written stimulus software based on StimGen (https://github.com/mscaudill/neuroGit; <u>Ruediger, 2020</u>) and presented on a gamma corrected LCD monitor (DELL, mean luminance: 60 cd/m2, monitor refresh rate 60 Hz: dimensions: 47.5 × 30 cm; 1680 × 1050 pixels). The monitor was positioned 15 cm from the left eye (contralateral to the right SC) and was angled at 60° from the long body axis. On the day of recording, mice were head fixed, and the Kwik-Cast protective covering was removed. Artificial cerebrospinal fluid (ACSF; 140 mM NaCl, 5 mM KCl, 10 mM d-glucose, 10 mM HEPES, 2 mM CaCl2, 2 mM MgSO4, pH 7.4) was used to keep the exposed tissue moist. Extracellular recordings of neural activity were collected using a linear silicon probe (Cambridge Neurotech, probe type: H3, 64-channel). The recording electrode was lowered into the brain using a micromanipulator (Luigs and Neumann) and stained with DiI (LifeTechnologies) for post-hoc identification of the recording site. Signals were amplified and band-passed filtered using 64 channels and Intan RHD headstage (Intan Technologies RHD2164) and recorded at 30KHz using an Intan RHD USB Interface Board. The tip of the electrode was lowered to 1700-1800uM below the pial surface to cover the superficial (visual) and intermediate levels of the SC. Recordings for control conditions started ~20 minutes after the probe was lowered to the target depth. Mice passively viewed a random sparse noise stimulus during which black squares covering 10° of visual space were presented at random locations of a 14×20 grid. Stimuli were presented on a white screen for a duration of100ms with a 200ms inter-stimulus interval. Stimuli were presented at each location for 20 repetitions. Without moving the electrode, prism lenses were gently placed and after 10 minutes the visual stimulus presentation was repeated

For UCLA Miniscope calcium imaging experiments, mice underwent two serial procedures spaced 4 weeks apart. During the first surgery, a 1 mm diameter craniotomy was made above the superior colliculus on the right hemisphere (centered at AP, −3.6 mm; ML, +0.7 mm from bregma). Then, 400 nl of AAV1-hSyn-Soma-GCaMP8m was injected into superior colliculus on the right hemisphere (AP, −3.6 mm; ML, +0.7 mm; DV, −1.7 mm). After the pipette was removed, the mouse remained on the stereotaxic frame for 20 min to allow complete diffusion of the virus. After 20 min of diffusion, the cortex below the craniotomy was aspirated with a 27-gauge blunt syringe needle attached to a vacuum pump, while constantly being irrigated with cortex buffer. When the striations of the corpus callosum were visible, the 27-gauge needle was replaced with a 30-gauge needle for finer-tuned aspiration. Once most of corpus callosum was removed, bleeding was controlled using surgical foam (Surgifoam), and then a 1 mm diameter × 4 mm length GRIN lens (GoFoton) was slowly lowered into the craniotomy. The lens was fixed with cyanoacrylate, and then dental acrylic was applied to cement the implant in place and cover the rest of the exposed skull. The top of the exposed lens was covered with Kwik-Sil (World Precision Instruments) to protect it and the Kwik-Sil was covered with dental cement.

Then, 4 weeks later, the mice were again put under anesthesia to attach the baseplate, visually guided by a miniscope. The overlying dental cement was drilled off and the Kwik-Sil was removed to reveal the top of the lens. A 3D printed headplate for head-fixation experiments was cemented. Then, the miniscope, with an attached baseplate, was lowered near the implanted lens and the field of view was monitored in real-time on a computer. The miniscope was rotated until a well-exposed field of view was observed, at which point the baseplate was fixed to the implant with cyanoacrylate and dental cement. The mouse did not receive post-operative drugs after this surgery as it was not invasive.

Mice were headfixed on a Styrofoam ball and the UCLA Miniscope Team’s Graphical User Interface (GUI) was used for acquiring, controlling, and visualizing Miniscope data. Visual stimuli were presented in Matlab with Psychtoolbox (methods described above). Stimuli consisted of a 2° by 2° black square moving at a rate of 30° per second along 60° of elevation (dorsal-to-ventral). The black square was presented at 13 different locations, spaced 10° apart, along the azimuth (nasal-temporal axis). After 6 stimulus presentations at each location, prism goggles (40° left shifting) were placed in front of the mouse and the visual stimuli were presented again.

Miniscope data was analyzed by selecting a region of interest (ROI) in the video corresponding to the calcium fluctuations tied to the visual stimulus position moving at the 0° nasal-temporal position of the mouse’s visual field. The raw pixel intensity across the ROI was then averaged at each timepoint and subtracted by an out-of-frame ROI for background fluctuations in the Miniscope image sensor. The max ROI response during each stimulus presentation was averaged for a given mouse to find the tuning curve for the ROI response.

#### Cortical lesions

Mice were anesthetized with 1.5-2% isoflurane and placed in a stereotactic set up (Kopf). Body temperature was monitored using a rectal probe and maintained at 37°C using a heating pad (FHC; DC Temperature Controller). Throughout the procedure, the animals’ eyes were protected by a thin layer of eye ointment (Bausch & Lomb). Animals had undergone goggle base implantation prior to behavioral training and so only the fur on the posterior half of the head was shaved. Skin was sterilized with Betadine before being removed to expose the skull. Lidocaine (7 mg/kg) was injected locally at the site of incision. The outline of primary visual cortex was traced on top of both hemispheres of the skull using permanent marker. For each hemisphere, a 0.07mm drill bit was used to thin along the marked outline until the piece of skull covering V1 was detached from the rest of the skull. The skull flap was gently lifted away and removed using a micro knife. The dura was then gently removed using the tip of an insulin needle and the micro knife was used to cut along the edges of the craniotomy to a depth of 1mm. A sharpened micro spoon was then used to remove the section of cortical tissue.

Surgifoam soaked in phosphate buffered saline was applied to the lesion site in between each step to reduce bleeding and maintain tissue integrity. Once lesions were complete on both hemispheres, the Surgifoam was removed and lesion sites were sealed with a local application of Kwik-Cast (WPI). The entire skull was then covered with dental cement (Ortho-Jet acrylic resin, Lang Dental). At the end of the procedure, animals were administered a subcutaneous injection of 0.1mg/kg Buprenorphine as a postoperative analgesic.

